# Production of a Highly Immunogenic Antigen from SARS-CoV-2 by Covalent Coupling of the Receptor Binding Domain of Spike Protein to a Multimeric Carrier

**DOI:** 10.1101/2021.04.25.441271

**Authors:** Argentinian AntiCovid Consortium, Paula M. Berguer, Matías Blaustein, Luis M. Bredeston, Patricio O. Craig, Cecilia D’Alessio, Fernanda Elias, Paola C. Farré, Natalia B. Fernández, Hernán G. Gentili, Yamila B. Gándola, Javier Gasulla, Gustavo E. Gudesblat, María G. Herrera, Lorena I. Ibañez, Tommy Idrovo-Hidalgo, Alejandro D. Nadra, Diego G. Noseda, Carlos H. Paván, María F. Pavan, María F. Pignataro, Ernesto A. Roman, Lucas A. M. Ruberto, Natalia Rubinstein, María V. Sanchez, Javier Santos, Diana E. Wetzler, Alicia M. Zelada

## Abstract

Since the discovery of SARS-CoV-2, several antigens have been proposed to be part of COVID-19 vaccines. The receptor binding domain (RBD) of Spike protein is one of the promising candidates to develop effective vaccines since it can induce potent neutralizing antibodies. We previously reported the production of RBD in *Pichia pastoris* and showed it is structurally identical to the protein produced in mammalian HEK-293T cells. In this work we designed an RBD multimer construct with the purpose of increasing RBD immunogenicity. We produced multimeric particles by a transpeptidation reaction between the RBD expressed in *P. pastoris* and Lumazine Synthase from *Brucella abortus* (BLS), which is a highly immunogenic and very stable decameric protein of 170 kDa. We vaccinated mice with two doses 30 days apart, and then we measured humoral immune response. When the number of RBD copies coupled to BLS was high (6-7 RBD molecules per BLS decamer, in average), the immune response was significantly better than that elicited by RBD alone or even by RBD-BLS comprising low number of RBD copies (1-2 RBD molecules per BLS decamer). Remarkably, the construct with high number of RBD copies induced high IgG titers with high neutralizing capacity. Furthermore, a superior immune response was observed when Al(OH)3 adjuvant was added to this formulation, exhibiting a higher titer of neutralizing antibodies. Altogether our results suggest that RBD covalent coupled to BLS forming a multimer-particle shows an advantageous architecture to the antigen-presentation to the immune system which enhances immune responses. This new antigen should be considered a potent candidate for a protein-based vaccine.

## Introduction

The coronavirus SARS-CoV-2, which was detected about one year ago in Wuhan, Republic of China in December 2019, has produced over 3.0 million deaths by April, 2021 (https://ourworldindata.org/covid-deaths). Since the discovery of the new strain, several vaccines have been developed (https://covid19.trackvaccines.org/vaccines/), based on inactivated viruses, DNA, RNA and/or proteins^1–3^. Furthermore, multivalent antigens that mimic antigen presentation by viral particles have been shown to induce a stronger immune response than monomeric antigens^4^, thus allowing immunization with lower doses of antigen, as well as the combination of different antigen variants. This approach would be critical in the specific case of new circulating SARS-CoV-2 lineages that have unfortunately emerged around the world^5^.

The receptor binding domain (RBD) of SARS-CoV-2 Spike protein is responsible for the interaction of the virus with the receptor Angiotensin-Converting Enzyme 2 (ACE2)^6^, which allows the penetration of the virus into the cells. Therefore, RBD constitutes a suitable target to develop diagnostic and therapeutic tools, as well as vaccines. It has been found that several neutralizing antibodies produced upon SARS-CoV-2 infection are directed towards RBD^7,8^. The structure of the complex formed by SARS-CoV-2 RBD and human ACE2 was recently determined^9^. Unlike SARS-CoV-1, several residue changes in SARS-CoV-2 RBD stabilize two virus-binding hotspots at the RBD/ACE2 interface increasing the affinity for ACE2^9^.

SARS-CoV-2 Spike’s RBD is a challenging protein to produce in heterologous systems because it has nine cysteine residues (eight of which form disulfide bonds), a complex topology and two N-glycosylation sites^10,11^. For these reasons, RBD is often expressed in mammalian cells^9,12^. Nevertheless, our AntiCovid consortium was able to produce RBD at high yields in *Pichia pastoris.* Its immunogenicity, stability and biophysical properties were similar to those of RBD produced in mammalian HEK-2913T cells; notably, combined with adjuvants it elicited humoral immune response in mice^12^. Remarkably, the production of RBD in yeast cells is more cost effective compared to other systems, and the fermentation bioprocess can be scaled up under Good Manufacturing Practice conditions. We were capable of optimizing RBD expression in *P. pastoris* to reach yields of 180 mg/L of ~90% pure protein, and thus were able to supply it to the Argentinian scientific community and health system at a low cost. It is currently used as an antigen for serological test kits, and as an immunogen. RBD could be used to produce neutralizing antibodies at large scale in animal systems such as egg yolk (IgY)^13,14^ or in horses^15^.

While RBD is immunogenic *per se*, the development of a highly immunogenic antigen is essential for the development of new generation subunit vaccines, which ideally should be capable of simultaneously providing immunity against the different emerging RBD variants. A possible strategy to achieve this goal is to couple the protein of interest to an immunogenic carrier protein. Lumazine synthase (LS, EC 2.5.1.78) from *Brucella abortus* (BLS) is a highly immunogenic and very stable decameric protein ideally suited to function as an immunogenic carrier protein. Five BLS subunits (17 kDa each) form a pentamer, and two pentamers form a very stable decamer. Such decamers may be decorated with any protein whose antigenicity needs to be increased. The BLS decamer has been used to significantly increase the immunogenicity of other proteins by means of the fusion between the C-terminus of the foreign protein and the N-terminus of BLS, showing that BLS is a convenient platform for antigen display^16–18^. Moreover, BLS activates dendritic cells *in vitro*, increasing the levels of co-stimulatory molecules and the secretion of proinflammatory cytokines, and also recruits dendritic cells *in vivo*, in both cases in a TLR4-dependent manner^19^. In addition, BLS chimeras are extremely effective to rapidly activate specific CD8+ lymphocytes, and to induce significant cytotoxic activity^20^. While other carrier proteins have been non-covalently attached to target proteins through a pair of interacting proteins (*e.g.* X-dockerin-cohesin-Y^21^ where X and Y are proteins expressed as a fusion protein with the dockerin and cohesin domains, respectively), such non-covalent interactions might be relatively weak particularly for immunization purposes, which normally require the use of strong adjuvants. The translational fusion of target proteins and peptides with BLS has allowed researchers to overcome this problem^22^. However, given that BLS fusions are usually expressed at high yields in *Escherichia coli*, this strategy may be not adequate for proteins with a complex structure and high disulfide bonds content, and/or that require post translational modifications, as it is the case of RBD, which cannot be properly expressed in *E. coli*^12^. This problem could be circumvented through the covalent coupling of RBD and BLS, *in vitro,* in their native state, which can be achieved with the Sortase A enzyme.

Sortase A and its variants can efficiently catalyze a transpeptidation reaction that creates a covalent link between two native proteins or peptides^23^. This reaction requires that the N-terminal protein to be covalently linked contains a specific signal located in its C-terminal stretch: **Protein**^**N**^-Leu-Pro-X-Thr-Gly **(Figure 1)**. This site should preferably be in an unstructured context, and the Gly residue is usually followed by a histidine tag to facilitate its purification process. The C-terminal protein to be covalently linked should contain three Gly amino acid residues in its N-terminal region (Gly-Gly-Gly-**Protein**^**C**^). After the Sortase A transpeptidation reaction, both subunits are joined to produce the covalently-linked product **Protein**^**N**^-Leu-Pro-X-Thr-Gly-Gly-Gly-**Protein**^**C**^ (**Figure 1**). Therefore, the Sortase A-mediated transpeptidation reaction can couple preassembled oligomeric carriers such as BLS, or virus-like particles (VLPs), to an immunogen expressed in a different system^24^. An additional advantage of Sortase A-mediated coupling is that the scar of the transpeptidation reaction in the protein product is of only seven residues, of which three are glycines.

**Figure 1.**
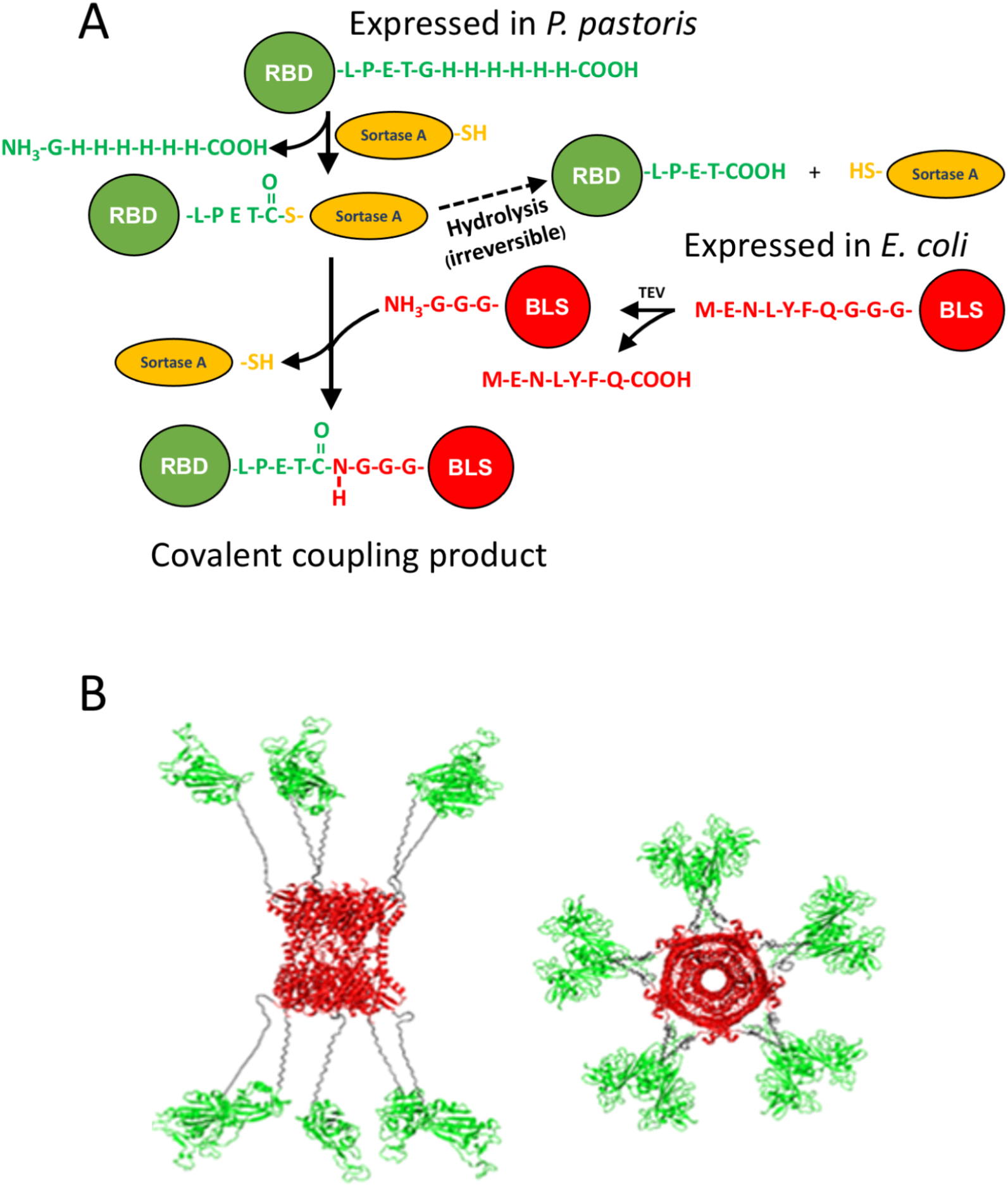
The Strategy: Sortase A-mediated Transpeptidation Reaction. (A) The reaction is exemplified by the coupling of a BLS subunit (red) and the SARS-Co-V2 RBD domain (green). The enzymatic intermediary form (RBD covalent bound to Sortase A) is shown, and the hydrolysis of the intermediary to yield the by-product **RBD**-Leu-Pro-Glu-Thr is included. The latter cannot be reused in the coupling reaction. The reaction catalyzed by TEV protease to yield Gly-Gly-Gly-BLS, the substrate of Sortase A is also shown. (B) A hypothetical reaction product with a multiplicity of 10:10 (RBD_10_/BLS_10_) is shown.

In this work, we produced a multimeric RBD-BLS by a Sortase A-mediated transpeptidation covalent coupling reaction between RBD expressed in *P. pastoris* and BLS produced in *E. coli*, with the purpose of increasing RBD immunogenicity. RBD-BLS induced a strong humoral response against SARS-CoV-2 RBD. Remarkably, the RBD antigen produced a significantly higher humoral response when presented to the immune system in the context of an RBD-BLS multimer with a high number of RBD molecules per particle (high multiplicity) than as a monomer, or within low multiplicity particles. The RBD-BLS multimer is a potential vaccine candidate, and can also be useful to produce neutralizing antibodies in animal systems. This platform could be also used to simultaneously present several proteins to the immune system, such as different RBD variants and other domains of the spike protein.

## Results

### Production of Sortase A-Substrates: BLS in E. coli and RBD in Pichia pastoris

In order to obtain a multimeric carrier for RBD, we first produced an oligomeric form of BLS in *E. coli*. For this purpose we synthesized and sequenced a gene encoding a BLS carrier protein variant which included in its N-terminal region a TEV protease cleavage site (in italics) followed by a Gly-tag (*Met-Glu-Asn-Leu-Tyr-Phe-Gln*-Gly-Gly-Gly-**BLS**) (**Figure 1**). This strategy allows the removal of the N-terminal Met after a quantitative digestion with the TEV protease, thus producing a polypeptide carrying an N-terminal Gly-Gly-Gly required for the Sortase A-mediated transpeptidation reaction. Following BLS expression in *E. coli*, the protein was purified in two steps, by anion exchange chromatography (Q-Sepharose) followed by size exclusion chromatography (SEC-HPLC, Superdex S200). The protein was obtained with a high yield (100 mg L^−1^, >98% purity), and its identity was verified after purification by mass spectrometry. BLS was then digested with TEV protease to generate the N-terminal Gly-Gly-Gly motif so that it can function as a Sortase A substrate. The digestion was confirmed by SDS-PAGE and mass spectrometry (**Figures S1 and S2**), which showed the expected “activated” multimeric carrier Gly-Gly-Gly-**BLS.** It is worth mentioning a complete TEV digestion of BLS was observed.

To generate the second substrate of the coupling reaction, an RBD variant including a Sortase A recognition signal (Leu-Pro-Glu-Thr-Gly-) followed by a (His)_6_-tag at its C-terminus was produced in *P. pastoris* (**RBD-***Leu-Pro-Glu-Thr-Gly*-(His)_6_) and obtained from culture supernatants with a high yield (180 mg L^−1^, >90% purity). We have previously shown that this protein is conformationally stable and drives a significant immune response in mice when administered in a formulation with adjuvants^25^. We assessed the tertiary structure of the purified RBD through near-UV-CD spectroscopy. The results are compatible with an asymmetric and rigid environment of the aromatic side chains, indicating a native packing (**Figures 2A and B**). Moreover, the protein does not exhibit significant aggregation, as inferred from the SEC-HPLC profiles and the absence of species with low elution volumes (**Figure 2C**). In addition, as previously observed^25^, SEC-HPLC analysis revealed the existence of two different species (differing in their glycosylation patterns) with elution times compatible with molecular masses between 45 kDa and 20 kDa.

**Figure 2.**
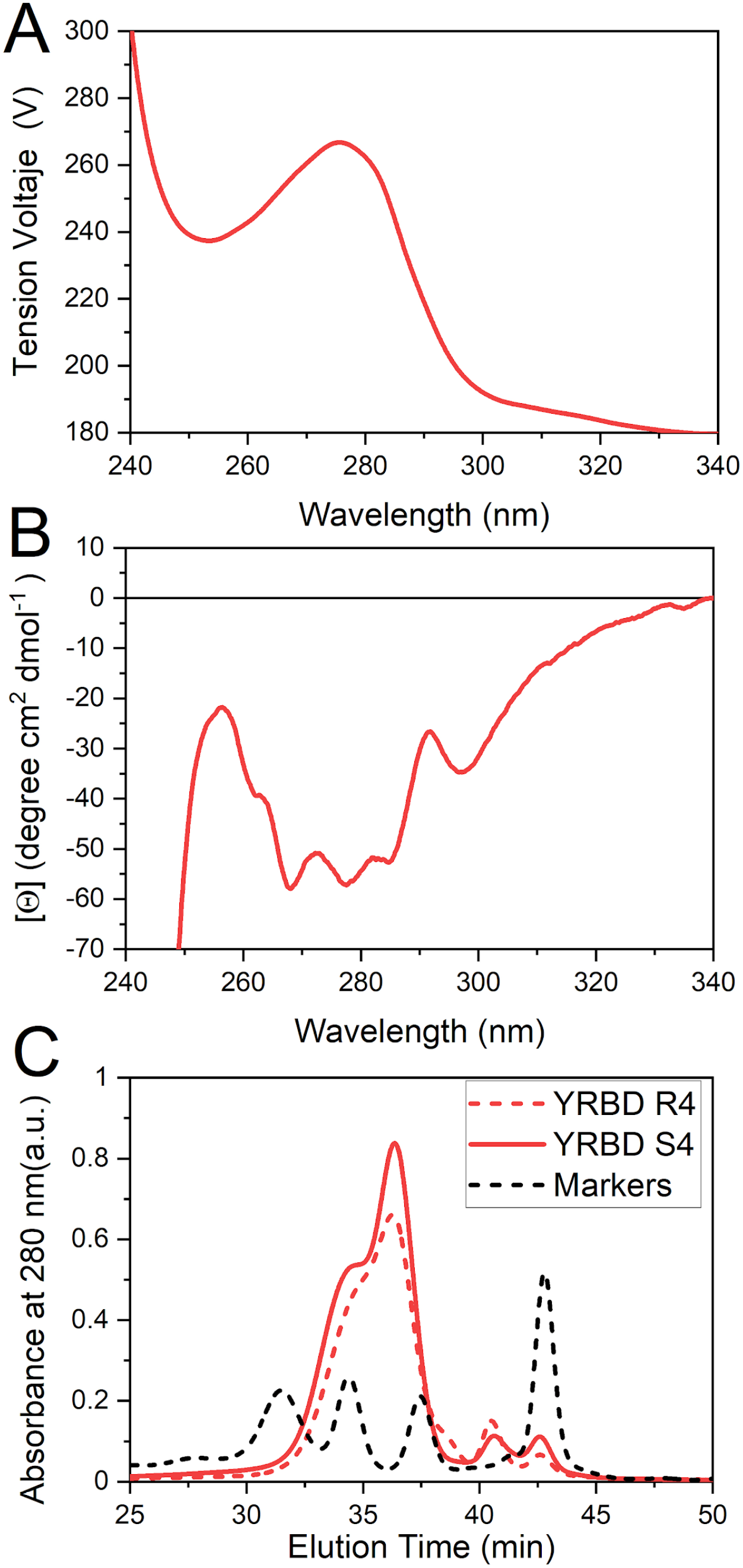
RBD Expressed in *Pichia pastoris* is Properly Folded and does not Aggregate. Near-UV Circular Dichroism Spectra of purified RBD: High tension voltage (A) and Near-UV CD spectra (B). The protein concentration was 30 *μ*M. The buffer used was 20 mM Tris-HCl, 100 mM NaCl, pH 7.4. (C) SEC-HPLC of RBD elution profiles (monitored at 280 nm) corresponding to two different batches (R4 and S4) of independently expressed and purified RBD, and of molecular mass markers (158, 44, 17 and 1.35 kDa).

### Native Sortase A Mediated Coupling of RBD to a Fluorescent Peptide

To evaluate the ability of recombinant **RBD**-*Leu-Pro-Glu-Thr-Gly*-(His)_6_ to act as a Sortase A substrate, we first carried out a preliminary reaction using the fluorescent peptide probe Gly-Gly-Gly-Ser-{Lys-**(TMR)**} as a substrate instead of BLS. This peptide contains a covalently bound tetramethylrhodamine (TMR) label. The product of this reaction was the covalently coupled **RBD**-Leu-Pro-Glu-Thr-Gly-Gly-Gly-Gly-Ser-(Lys-**TMR**) (RBD-TMR).

After the coupling reaction and SEC (G25) separation, nearly 17% of input RBD was labeled with the fluorescent probe as inferred from UV and visible spectra analysis (**Figures 3A and B)**. The unreacted RBD retained the (His)6 tag, and could therefore be separated and recovered by addition of NTA resin to the reaction mix followed by centrifugation. This procedure produced a final yield of ~50% labelled RBD. Moreover, SEC-HPLC analysis, monitored at 554 nm, which allows detection of the TMR label, revealed the presence of the reaction product. However, no free fluorescent peptide was observed in the elution profile (**Figure 3C)**, indicating that the G25 matrix was capable of separating the residual free peptide. SEC-HPLC (using the Superose 6 column) of the reaction product showed an absence of peaks at short elution times, suggesting that RBD-TMR was not prone to aggregation after the covalent coupling of the probe. These results showed that Sortase A was functional, and it was possible to carry out the covalent coupling between recombinant RBD and a Gly-Gly-Gly-tagged substrate.

**Figure 3.**
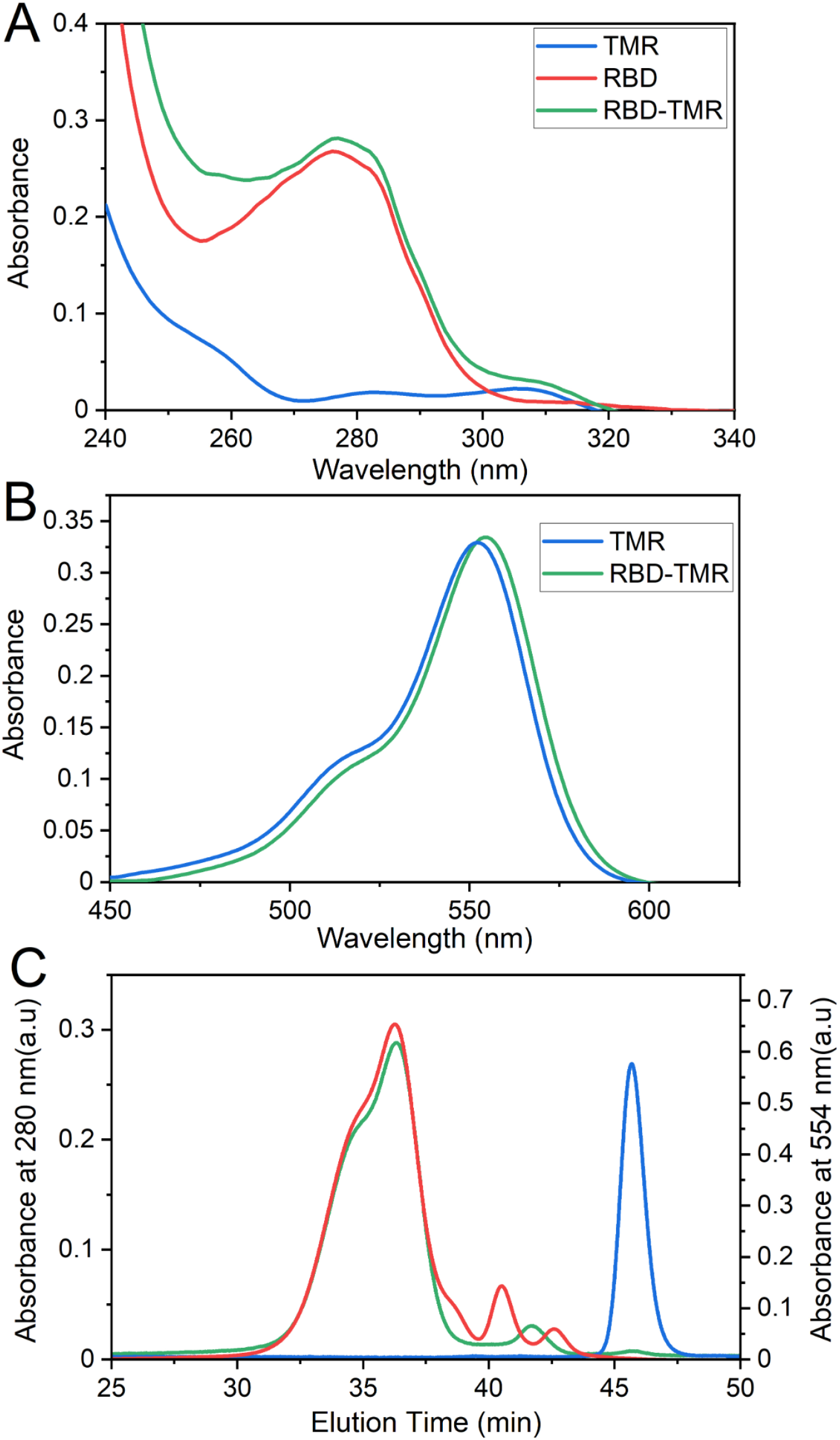
Covalent Coupling of RBD to a Fluorescent Probe Mediated by Sortase A. (A) UV and (B) visible spectra of the free fluorescent probe (blue) and the coupling reaction product RBD-TMR (green) after a G25 column separation. The RBD spectrum is also shown as a reference (red) (C) SEC-HPLC elution profiles (Superose 6 column) of RBD (monitored at 280 nm, red), TMR (monitored at 554 nm, blue) and RBD-TMR (monitored at 554 nm, green) after coupling and G25 separation.

### Functionality of Recombinant RBD

In addition to the conformational analysis, we also assessed the functionality of purified RBD-TMR. Therefore, we tested its ability to bind RBD natural receptor during viral infection, the human ACE2. For this purpose, we incubated (a) the labeled protein (RBD-TMR), (b) a soluble version of ACE2 with a C-terminal (His)6 tag, and (c) agarose-NTA-Ni^2+^resin **(Figure 4A**) during 30 min. Next, we separated the ternary complex ACE2/RBD-TMR/agarose-NTA-Ni^2+^ by centrifugation. The matrix agarose-NTA-Ni^2+^ interacts with the (His)_6_ tag that it is present only in the C-terminal of ACE2 (and not in RBD-TMR), thus pulling down bound RBD-TMR and consequently producing a significant decrease of the supernatant fluorescence intensity (570-580 nm, **Figure 4B**). Elution of the complex from the NTA matrix by addition of 300 mM imidazole caused a strong fluorescence increase in the soluble fraction, further indicating that (His)_6_-tagged ACE2 forms a complex with RBD-TMR (**Figure 4C**). We concluded that RBD-TMR was capable of interacting with the ACE2 soluble domain, indicating that the structural features relevant for the interaction are preserved in recombinant RBD produced in *P. pastoris*, and that the conditions employed to perform the transpeptidation reaction to produce RBD-TMR did not affect its tertiary structure.

**Figure 4.**
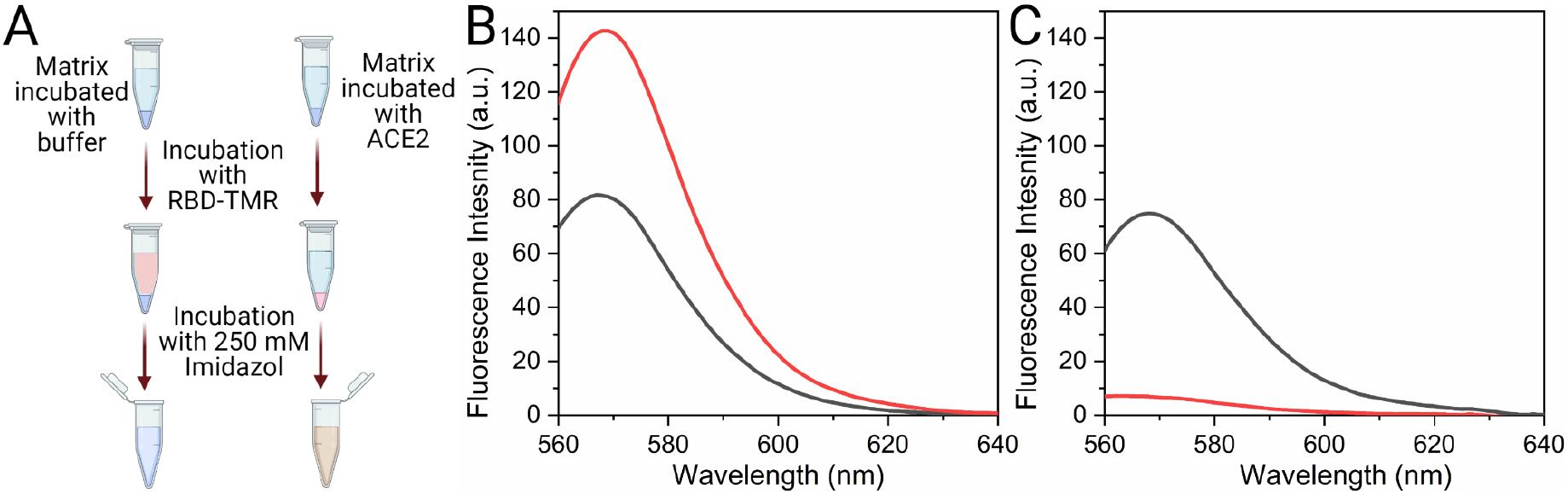
Evaluation of RBD-TMR Binding to ACE2. (A) RBD-TMR pull down and elution from the matrix. (His)_6_-tagged ACE2 was immobilized in an NTA matrix (300 μL of 0.9 μM ACE2 + 75 μL of agarose-NTA-Ni^2+^resin). A control incubation was performed omitting ACE2. Samples were centrifuged and washed three times and subsequently incubated in the presence of 1 μM RBD-TMR for 30 min under mild agitation. After centrifugation, the supernatant was transferred to a fresh tube and analyzed by fluorescence spectroscopy (B). The matrix was washed three times and eluted with 300 μL of 20 mM Tris-HCl pH 8.0, 150 mM NaCl and 250 mM imidazole. The sample was centrifuged, and the supernatant was analyzed by fluorescence spectroscopy (C). In B and C black lines correspond to the incubation in the presence of ACE2, while red lines correspond to the controls without ACE2. The excitation wavelength was 550 nm, the bandwidths for both excitation and emission were of 5 nm. All the measurements were made at 25 °C.

### Native Sortase A-mediated Coupling of RBD to BLS

Given that recombinant RBD proved to be a good Sortase A substrate, we proceeded to attempt the coupling of **RBD**-Leu-Pro-Glu-Thr-Gly-(His)_6_ and TEV-digested Met-Glu-Asn-Leu-Tyr-Phe-Gln-Gly-Gly-Gly-BLS. The expected product was the covalently coupled **RBD**-Leu-Pro-Glu-Thr-Gly-Gly-Gly-Gly-Ser-Gly-Ser-Gly-**BLS**(RBD-BLS in **Figure 1**). The SEC-HPLC profiles (Superose 6 column) corresponding to a BLS multimer, RBD alone and the reaction product are shown in **Figure 5A**. The product of the covalent coupling mediated by Sortase A exhibited an elution time compatible with an increment in the hydrodynamic radius of the assembly compared to that observed for the decameric BLS (27 and 31.5 min, respectively). The unreacted BLS decamer and RBD are also clearly distinguishable in the SEC profile.

**Figure. 5.**
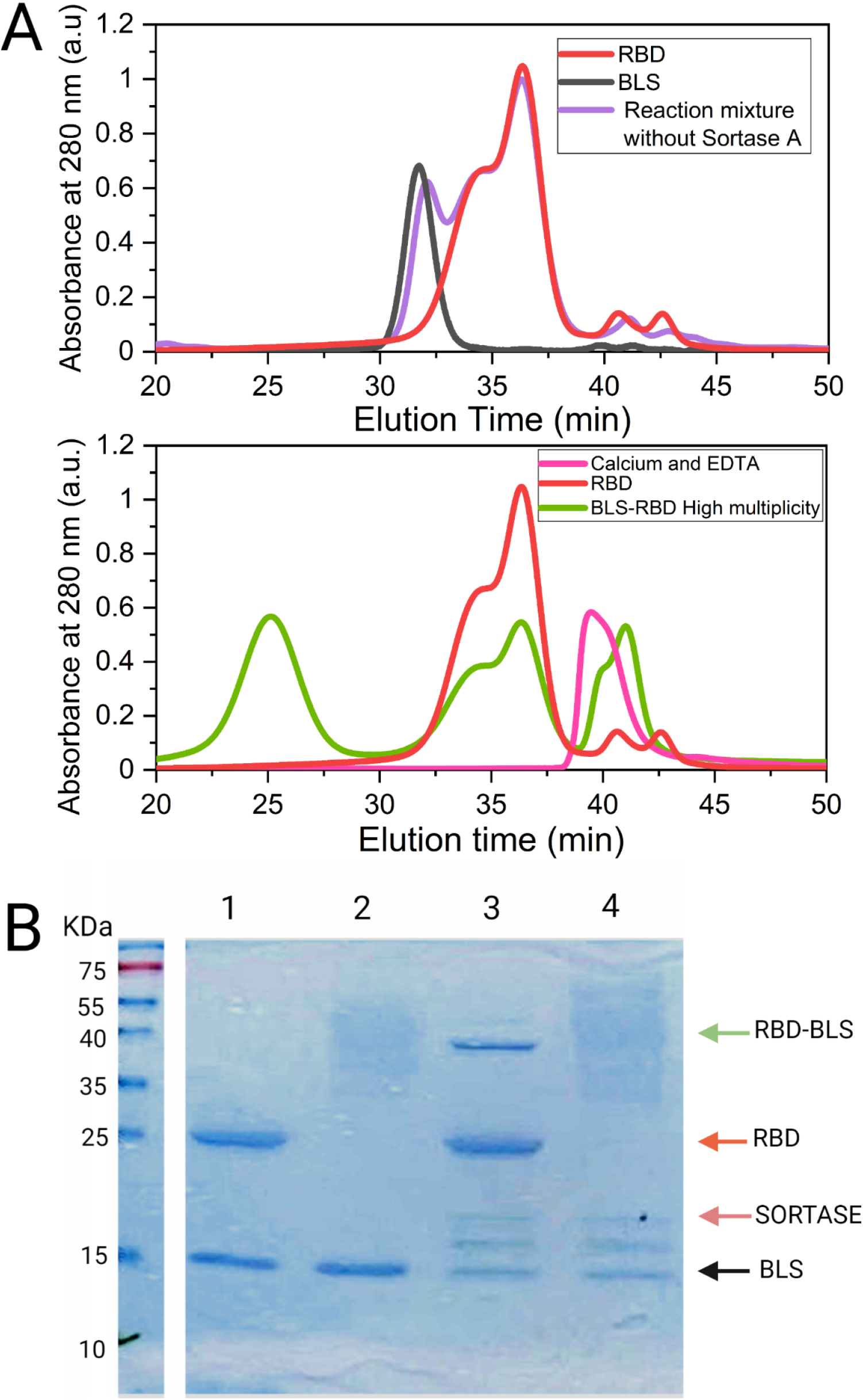
Sortase A-mediated Covalent Coupling of RBD and BLS. (A) SEC-HPLC profiles corresponding to Gly-Gly-Gly-BLS (black), RBD (red), the control reaction mixture without Sortase A (violet) and the product of the reaction catalyzed by Sortase A, (green, lower panel). An equivalent volume of a 10 mM EDTA, 10 mM CaCl_2_ solution was loaded as a control (magenta, lower pane). Decameric BLS has a molecular weight of approximately 170 kDa and each RBD subunit adds approximately 26 kDa or 40 kDa (excluding or including glycosylation, respectively). The observed multiplicity was ~6.6 (RBD_6.6_/BLS_10_) with a yield of ~20%. The RBD-BLS oligomer eluted at ~25 min. The peak observed between 38-43 min corresponds to EDTA-Ca^2+^. Sortase A requires 10 mM CaCl_2_ for the activation of its catalytic activity, therefore 10 mM EDTA was added to stop the reaction, which was carried out during 240 min at 4 °C. (B) SDS-PAGE analysis of Sortase A-mediated RBD-BLS coupling. To facilitate the detection of the Sortase A reaction products, the protein samples were treated or not with endoglycosidase H. Lanes 1-2: RBD + BLS without Sortase A, deglycosylated (lane 1), or not (lane 2), Lanes 3-4: RBD+BLS after reaction with Sortase A, deglycosylated (lane 3) or not (lane 4).

Since BLS is a decameric protein, different multiplicities (RBD_n_/BLS_10_) can be obtained depending on the buffer conditions and the stoichiometry of the Sortase A reaction. It was possible to obtain different multiplicities by controlling RBD and BLS concentration ratios. We used low RBD and high BLS concentration, to obtain low multiplicities (between 1 and 2 of RBD copies per multimer, **Figure S3**), referred as RBD-BLS LM), whereas the higher multiplicities (referred as RBD-BLS HM) were obtained with higher RBD/BLS ratios (**Figure S3** and **Figure 5A**).

To determine such multiplicities, and given that the presence of heterogeneous sizes of glycan moieties in RBD resulted in a high heterogeneity which difficults the analysis, we trimmed the glycans with endoglycosidase H and resolved the deglycosylated products of the coupling reaction by SDS-PAGE (**Figure 5B**, compare lanes 1 and 3 to 2 and 4). The sharp band that migrates close to the 40 kDa marker in lane 3 corresponds to the covalently coupled RBD-BLS (26 kDa of deglycosylated RBD +17 kDa of monomeric BLS, 43 kDa, Lane 1). It should be noted that a large amount of RBD still remains uncoupled (blue arrow in Lane 3), therefore the efficiency of the reaction still needs to be improved. Additionally, a byproduct (between 15-18 kDa) was observed that still needs to be identified (lanes 3 and 4).

A quantification of the loss of free BLS and the increment of the RBD-BLS band in SDS-PAGE (and HPLC) allowed us to quantify the yield and the multiplicity (**Figure S4)**, that is the number of RBD copies per RBD-BLS particle, of the coupling reaction. We have obtained preliminary yields of 10-30% with average multiplicities between 2 and 4 (RBD_2_/BLS_10_ and RBD_4_/BLS_10_, respectively) in preliminary experiments. However, the incorporation of a concentration step (the reaction made in a centrifugal filter unit (Centricon, Merck Millipore) with a cutoff of 10 kDa, see Materials and Methods for details) allowed us to enhance the reaction, yielding in higher multiplicities (between RBD_6._/BLS_10_ and RBD_7._/BLS_10_, RBD-BLS HM).

In summary, we succeeded in coupling the eukaryotic-produced RBD antigen with the *E.coli* expressed decameric carrier BLS under native conditions. In addition, the possibility of obtaining RBD-BLS with different multiplicities allowed to study the effect of multiplicity on the development of the immune response.

Once purified by SEC, the coupling product was found to be stable after freezing and subsequent thawing, and did not show any significant aggregation tendency. In addition, the product could be easily concentrated, and did not show any apparent interaction with the concentrator membranes (data not shown). Importantly, the coupling products were also separated from the Sortase A enzyme by SEC. This is an essential step to avoid reverse transpeptidation and RBD-Sortase A hydrolysis (that would result in a RBDh product that can not be recycled by the enzyme)^26^, which would reduce the global yield of the reaction.

### The Immunogenicity of RBD is Increased by Coupling to the BLS multimer

To assess whether RBD coupled to BLS can improve the immunogenicity of RBD, we immunized mice with two doses of RBD-BLS HM, RBD-BLS LM and monomeric RBD, 30 days apart (**Table 1**, Materials and Methods). In addition, RBD and RBD-BLS-HM formulations were adjuvanted with Al(OH)_3_.

**Table 1.** Immunization Treatments.

| Groups | RBD<br>(µg/dose) | BLS<br>(µg/dose) | Covalent<br>coupling | RBD: BLS<br>(molar ratio) | Adjuvant<br>(Al(OH)<sub>3</sub>) |
| --- | --- | --- | --- | --- | --- |
| **RBD** | 15 | 0 | --- | --- | No |
| **RBD** | 15 | 0 | --- | --- | Yes |
| **RBD + BLS** | 15 | 17 | No | 6.6:10 | No |
| **RBD-BLS LM** | 15 | 70 | Yes | 1.5:10 | No |
| **RBD-BLS HM** | 15 | 17 | Yes | 6.6:10 | No |
| **RBD-BLS HM** | 15 | 17 | Yes | 6.6:10 | Yes |

The multimeric RBD-BLS HM formulation alone elicited a strong humoral response with high anti-RBD IgG titers, and when adjuvanted with AL(OH)_3_ the humoral response increased 6.75 times relative to RBD-BLS HM alone (**Figure 6**). By contrast, monomeric RBD did not elicit a significant anti-RBD IgG specific antibody response even in presence of Al(OH)_3_. We also observed that the addition of BLS to monomeric RBD (at the same BLS concentration used in the case of RBD-BLS HM) slightly improved humoral immune response (RBD + BLS group).This result suggests that BLS showed some adjuvant ability *per se*. Furthermore, the multiplicity of RBD-BLS multimers turned out to be a key variable, since a high multiplicity elicited strong immune response, while a low multiplicity did not (RBD-BLS LM group, between 1 and 2 copies of RBD per multimer).

**Figure 6.**
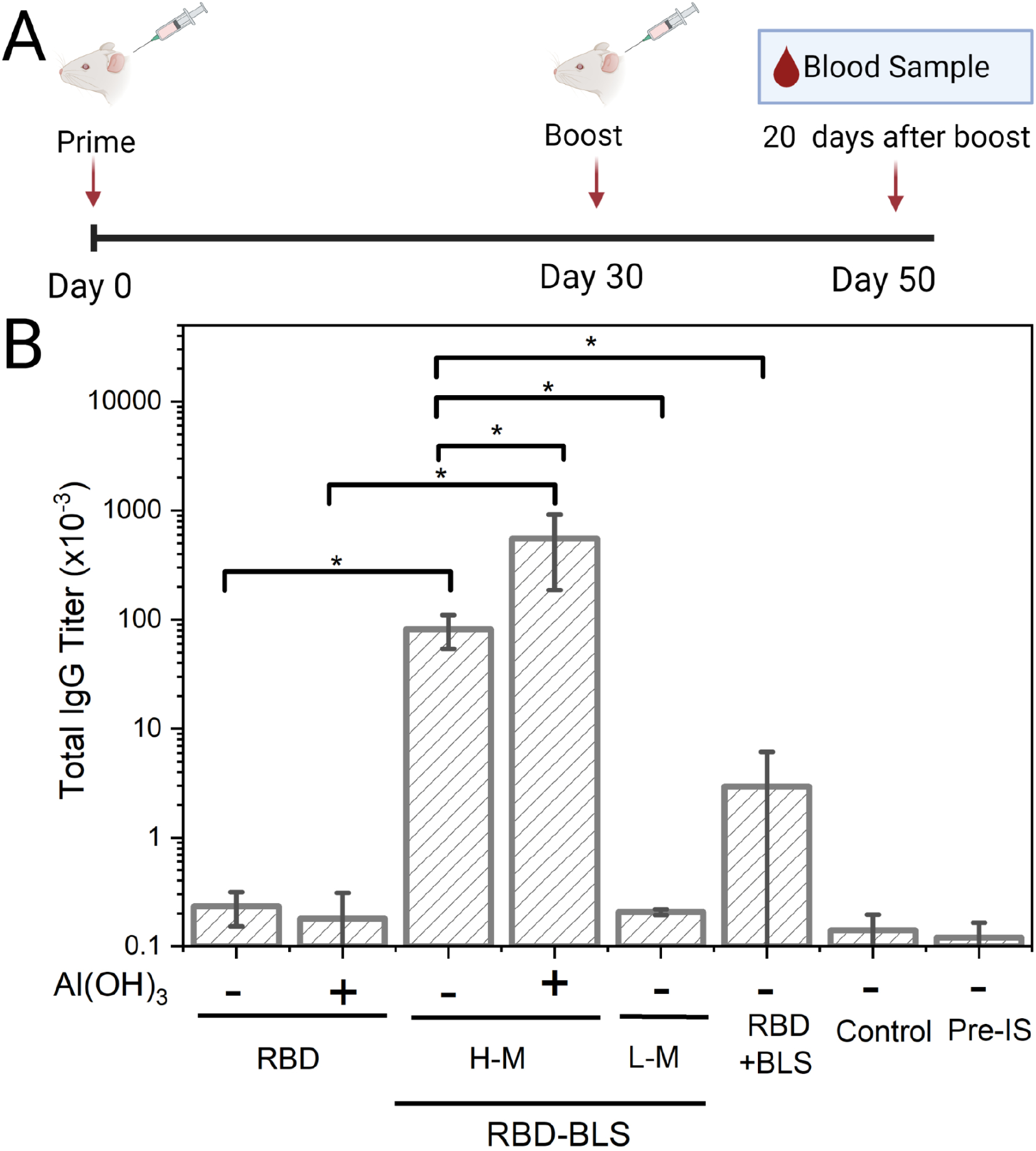
RBD-specific Humoral Immune Response in Mice Vaccinated with Monomeric RBD and RBD-BLS Multimers. A) Immunization scheme and blood collection. BALB/c mice (n=5) were vaccinated with two doses of the formulations 30 days apart subcutaneously and sera were obtained 20 days after boost. B) Evaluation of anti-RBD IgG antibody titers. Serial dilutions of sera samples were dispensed in a multi-well plate and RBD-specific IgG titers were determined by ELISA at day 50. Multiplicity (RBD/BLS), presence (500 μg) or absence of Al(OH)_3_ adjuvant, and whether or not BLS is covalently bound to RBD is indicated. The results are expressed as mean endpoint titers. The S.E.M. is indicated by vertical lines. Differences were statistically significant at p<0.05 (*).

These results clearly show that coupling of RBD to BLS at high multiplicity strongly enhances RBD immunogenicity, which might be due at least in part to the spatial arrangement of the RBD subunits in the resulting particles (**Figure 1B**).

### Immunization with BLS-RBD Elicit Antibodies with Neutralizing Activity Against a SARS-CoV-2 S Pseudotyped Virus Transduction

To analyze the neutralizing activity of immune sera elicited by RBD-BLS immunization, *in vitro* neutralization assays using SARS-CoV-2 S pseudotyped lentiviruses were performed^27^. Serial dilutions (1:100; 1:500; 1:2,500; 1:12,500; 1:62,500; 1:312,500; 1:1,562,500; 1:7,812,500) of sera were made in the DMEM medium and incubated with equal amounts of pseudotyped virus, then added to HEK-293T cells previously transfected with ACE2 and TMPRSS2 protease. While pseudotyped virus particles were capable of delivering GFP-encoding RNA into the cells in the presence of preimmune serum, immune sera prevented transduction, shown as a percentage of inhibition, in a concentration-dependent manner. Sera from mice immunized with RBD-BLS HM (**Table 1**), strongly reduced the number of GFP-expressing cells (**Figure 7**). Remarkably, immunizations with RBD-BLS HM administered in the presence of Al(OH)_3_ adjuvant resulted in significantly higher neutralizing antibody titers (**Figure 7**, IC_50_ values are 8700 and 2132, with and without adjuvant, respectively), as judged by the significant reduction observed in pseudotyped virus transduction. By contrast, sera from mice immunized with *Pichia pastoris*-expressed RBD domain plus BLS (not covalently coupled) resulted in lower neutralizing antibody titers (IC_50_ = 356.2, **Table 1**, RBD + BLS).

**Figure 7.**
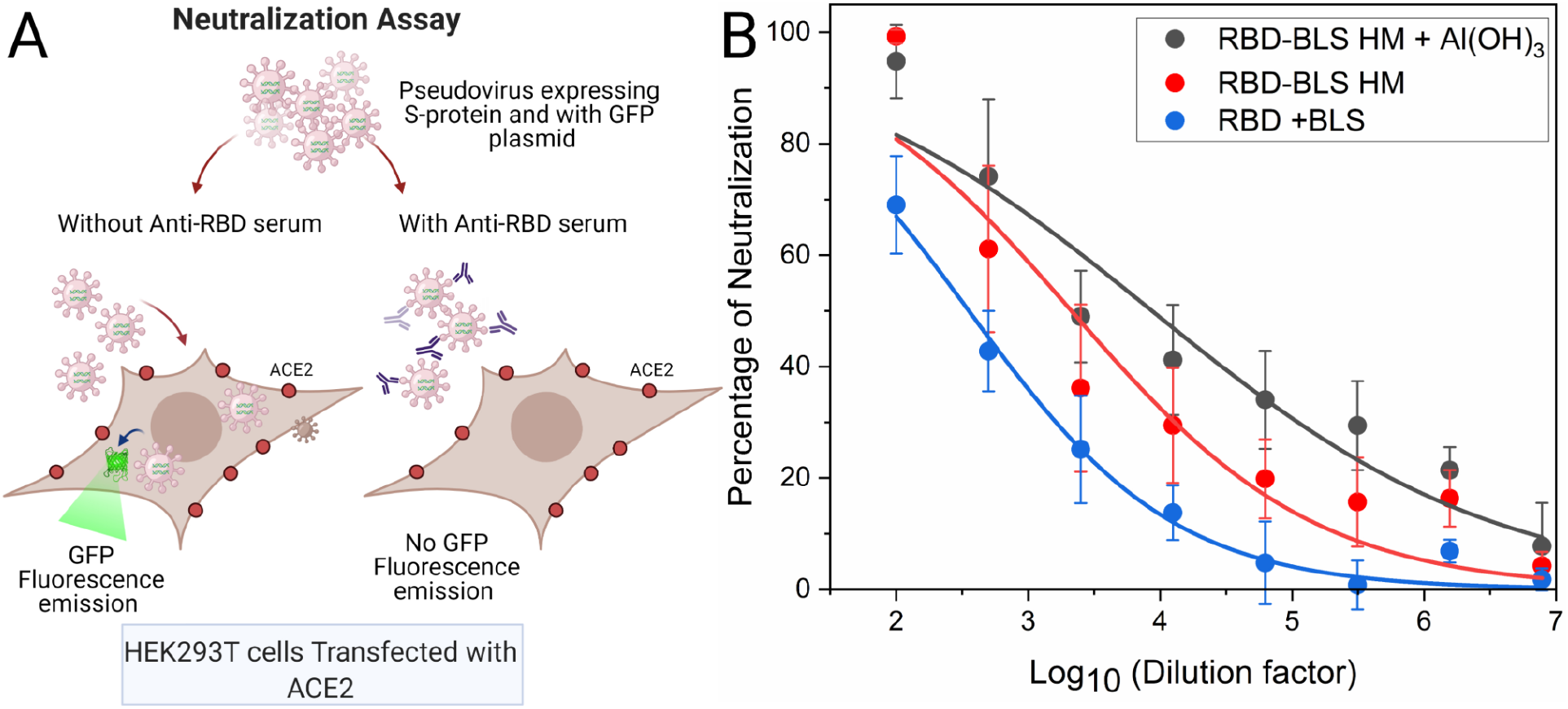
SARS-CoV-2 S Pseudotyped Virus Neutralization Assay. HEK-293T cells transfected with ACE2 and TMPRSS2 protease were transduced with aSARS-CoV-2 S pseudotyped lentivirus carrying a GFP-encoding mRNA in the presence of different dilutions of mouse sera. Forty eight hours later, cells were observed under the microscope. The number of GFP positive cells for each serial dilution was relativized to normal-serum control. The serum antibody dilution causing a 50% reduction of GFP positive cells (IC_50_) compared to control “virus only” treated cells was calculated using Graphpad Prism.

Altogether these results show that RBD-BLS HM is a very promising candidate both to induce neutralizing antibodies in animal models, and as a candidate for a protein-based vaccine.

## Discussion

A year ago, just before SARS-CoV-2 was first detected in Argentina, a group of researchers from different institutions and with different relevant expertises joined forces to help fighting the ongoing pandemic. Our work aimed to produce the RBD domain of SARS-CoV-2 spike protein, as it is a useful tool for the serological detection of infected patients, and has the potential to be used as antigen to develop a low cost vaccine. So far we have successfully produced RBD in different expression systems, among them in the yeast *P. pastoris,* which showed biophysical properties similar to that of RBD expressed in mammalian cells^25^.

In this work RBD was covalently coupled to a multimeric carrier with the aim of improving its immunogenicity to develop a useful vaccine candidate. As a carrier we used the multimeric lumazine synthase protein from *Brucella abortus* (BLS), which has been previously shown to be a good antigen carrier for systemic immunization and as mucosal-adjuvant, which makes it a good candidate to be used for oral subunit vaccines^17^. We have improved the efficiency of the RBD-BLS coupling reaction to scale up the production and purification of the RBD-BLS oligomer, obtaining different multiplicities (1 to 6) and we have compared the immunogenicity of RBD-BLS *vs*. RBD in mice.

Our results showed that the RBD-BLS multimer can elicit an increased immune response compared to that of monomeric RBD, even in the absence of adjuvant. This increase depended on the covalent coupling of RBD to BLS, as the co-administration of RBD with BLS (*i.e.* not covalently coupled) did not induce an equivalent immune response. We also observed that the multiplicity of RBD coupled to BLS was an important factor in the enhancement of the immunological response.

In immunological studies we observed that RBD coupled to BLS at a high multiplicity (6-7 RBD copies per BLS decamer) is remarkably more immunogenic than monomeric RBD, or than RBD coupled to BLS at a low multiplicity (1-2 RBD copies per BLS decamer). An 80-fold increment in the anti-RBD humoral immune response was observed by RBD-BLS HM compared to monomeric RBD. By contrast, no significant humoral immune response was elicited by monomeric RBD in the presence or absence of Al(OH)_3_, or by the addition of BLS to RBD (without covalent coupling). In addition, when Al(OH)_3_ was added to RBD-BLS HM an extra 6.75-fold increase in total anti-RBD IgG titer and 4.1 fold increase in sera neutralization IC_50_ value was observed.

Whereas BLS is highly resistant to urea-induced unfolding, the decameric structure is sensitive to low pH (pH 4.0-5.5), or to moderate guanidinium chloride concentrations (2.0 M)^28^, which cause the dissociation of the decameric forms to pentamers. In addition, BLS has been previously modified, and several pentameric variants also exist that could be used to assemble pentamers, which could then be mixed to obtain the decameric form, taking advantage of surface complementation that depends on the new side-chain residues on these surfaces^29^. This makes BLS a very malleable protein, and would allow, for example, to include different immunogens in the same protein particle. BLS oligomers carrying different coupled proteins could be disassembled by incubation at low pH, mixed and reassembled so as to generate a diversity of homo and hetero decamers. As a Toll-like receptor (TLR) agonist, it has been shown that BLS can stimulate both innate and adaptive immune responses, thereby improving vaccine efficacy^17,20^. Thus RBD-BLS, alone or in combination with adjuvants, has the potential to boost T- and B-cell responses against SARS-CoV-2, which deserves further exploration.

The idea of coupling RBD to a multimeric protein to produce a stronger immunogen has been considered by other groups. Some molecular platforms have been proposed to multimerize CoV antigens. A very interesting recent work reports the production of particles displaying regions of the Middle East respiratory syndrome coronavirus (MERS-CoV) spike protein, by coupling its RBD domain to a multimeric lumazine synthase (LS), using the SpyTag/SpyCatcher system^30^, in which the peptide SpyTag that includes 13 amino acids reacts with the 12.3 kDa-SpyCatcher protein^31^. A similar strategy has been recently developed by Hang and coworkers for SARS-CoV-2 spike protein using both *Aquifex aeolicus* LS and *Helicobacter pylori* ferritin as nanoparticle scaffolds^32^. They coupled a trimeric version of the antigen to LS, which caused such nanoparticles to elicit significantly higher neutralizing antibody responses than the spike protein alone. New strategies to develop novel vaccines against SARS-CoV2 also include self-assembling fusion proteins. For example, Powell and coworkers have recently prepared in Expi293F mammalian cells a fusion protein containing the spike ectodomain followed by *H. pylori* ferritin, a 19 kDa protein that self-assembles into a 24-subunit nanoparticle^33^. Of note, such nanoparticles display eight copies of a trimeric antigen on the surface, as previously used for haemagglutinin from H1N1 virus^34^. The authors showed that multivalent presentation of spike trimers on ferritin can notably increase elicitation of neutralizing antibodies in mice after a single dose.

In this context, the strategy presented here involving RBD-BLS particles may be considered as a new useful tool against SARS-CoV-2. The platform is ductile enough to incorporate new variants like E484K, which has been associated with escape from neutralizing antibodies^35–37^ or N501Y, which is associated with increased binding specificity and faster-growing lineages^38^. This is important given the high chance of having new waves of SARS-CoV-2 involving RBD mutant viruses.

## Materials and Methods

### SARS-CoV-2 RBD Protein Expression and Purification

The receptor binding domain of spike protein from SARS-CoV-2 (RBD) was expressed in *Pichia pastoris*. The RBD coding sequence (spike amino acid residues 319-537) was fused to the *Saccharomyces cerevisiae* alpha factor secretion signal (N-terminal) followed by a C-terminal Sortase A recognition sequence and a His_6_ tag (C-terminal) given the C-terminal sequence **RBD-**Leu-Pro-Glu-Thr-Gly-(His)_6_ as previously reported^25^. The expression was carried out in a bioreactor as indicated, and the protein was purified with a yield of 150 mg L^−1^ of culture supernatant^25^. RBD concentration was determined by UV spectrophotometry using an absorption coefficient of 33850 M^−1^ cm^−1^.

### BLS Expression and Purification

A gene encoding a variant of lumazine synthase of *Brucella abortus* (BLS) including an N-terminal TEV site followed by a Gly-Gly-Gly sequence and a Gly-Ser-GLy-Ser-GLy spacer (Met-Glu-Asn-Leu-Tyr-Phe-Gln-Gly-Gly-Gly-Gly-Ser-GLy-Ser-Gly-**BLS**) was assembled and cloned in the pET11a vector (Novagen). The construct was checked by sequencing. The protein was expressed in *E. coli* BL21(DE3). Cells were grown in Luria Bertani broth at 37 °C to OD_600_ 1.0, and protein expression was induced by 1 mM IPTG. Expression was carried out at 28 °C overnight. The cells were harvested by centrifugation at 6,000 rpm for 20 min and freezed at −20°C until use. The cell pellet was resuspended in 20 mL of 20 mM Tris-HCl, pH 7.4 and gently sonicated in an ice-water bath. The sample was centrifuged at 10,000 rpm for 30 min, and the soluble fraction was transferred to a fresh tube. The protein was purified in two steps. First the soluble fraction was loaded onto an anionic exchange chromatographic column (Q-Sepharose, given that the theoretical pI value of BLS is 6.4). The fractions containing BLS, as judged by SDS-PAGE analysis, were pooled and loaded onto a size exclusion chromatographic column (Superdex S200). Fractions containing BLS were pooled and the purified protein was analyzed by mass spectrometry and SDS-PAGE. BLS concentration was determined by UV spectrophotometry using an absorption coefficient of 18405 M^−1^ cm^−1^.

### ACE2 Expression and Purification

The expression plasmid pcDNA3-sACE2(WT)-8his, which includes the human ACE2 soluble domain (residues 1-615) coding gene, followed by a Gly-Ser-Gly linker and eight His residues for purification, was transfected in HEK-293T mammalian cells. Cells were grown in high glucose (4.5 g L^−1^) Dulbecco’s modified Eagle’s medium (DMEM, Thermo Fisher Scientific) supplemented with 10% fetal bovine serum (FBS, Natocor), 110 mg L^−1^ of sodium pyruvate (Thermo Fisher Scientific) and penicillin/streptomycin (100 units mL^−1^ and 100 μg mL^−1^ respectively, Thermo Fisher Scientific) in a 37 °C incubator under 5% CO_2_. Cells were plated (2×10^7^ cells per 150 mm plate) and grown for 24 h before transfection with Polyethylenimine (PEI, Sigma). After 72 h, the cell culture medium was centrifuged twice at 12,000 ×g for 20 min at 4 °C. The supernatant pH was adjusted to pH 8.0 with an equilibration buffer (50 mM sodium phosphate, 300 mM NaCl and 20 mM imidazole, pH 8.0). Soluble ACE2 was purified using a previously equilibrated Ni ^2+^-NTA-agarose column. The protein was eluted by increasing imidazole concentrations (prepared in the equilibration buffer). Fractions containing the ACE2, as judged by the SDS-PAGE analysis, were pooled and dialyzed against a buffer without imidazole (20 mM sodium phosphate, 150 mM NaCl, pH 7.4). pcDNA3-sACE2(WT)-8his was a gift from Erik Procko (Addgene plasmid # 149268; http://n2t.net/addgene:149268; RRID:Addgene_149268)^39^, and the HEK-293T mammalian cell line was kindly provided by Xavier Saelens (VIB-University of Ghent, Belgium). ACE2 concentration was determined by UV spectrophotometry using an absorption coefficient of 157220 M^−1^ cm^−1^.

### TEV Protease Expression and Purification

Tobacco Etch Virus protease (TEV) was a gift from Helena Berglund (Addgene plasmid # 125194; http://n2t.net/addgene:125194; RRID:Addgene_125194)^40^. Briefly, *E coli* Rosetta (DE3) cells (starting with a 1/20 dilution of an overnight pre-culture) were grown in Luria Bertani media at 37 °C up to OD_600nm_ of 0.6. TEV was induced (for 4 h at 37 °C) by the addition of 0.5 mM IPTG. Cells were centrifuged, resuspended, and sonicated in 20 mL of lysis buffer (50 mM phosphate pH 8.0, 150 mM NaCl, 20 mM Imidazole, 10% glycerol). The sample was subsequently centrifuged, and the soluble fraction loaded in a NTA-Ni^2+^ agarose column previously equilibrated with the lysis buffer. A washing step with 10 column volumes was performed, and the protein was eluted with 2.5 volumes of elution buffer (50 mM phosphate pH 8.0, 150 mM NaCl, 250 mM Imidazole, 10% glycerol). Finally, to remove imidazole, the protein was subjected to dialysis and stored at −80 °C in small fractions.

### Sortase A Expression and Purification

The Sortase A pentamutant (eSrtA) in pET29 was a gift from David Liu (Addgene plasmid # 75144; http://n2t.net/addgene:75144; RRID:Addgene_75144)^41^. Briefly, *E. coli* BL21 (DE3) cells (starting with a 1/40 dilution of an overnight pre-culture) were grown in Terrific Broth at 37 °C up to OD_600nm_ of 0.6. Sortase A expression was induced for 5 h at 37 °C by the addition of 1 mM IPTG. Cells were centrifuged, resuspended, and sonicated in 35 mL of lysis buffer (50 mM Tris-HCl pH 8.0, 150 mM NaCl, 20 mM Imidazole, 10% glycerol). After that the sample was centrifuged and the soluble fraction loaded in NTA-Ni^2+^ agarose previously equilibrated with the lysis buffer. A washing step with 10 column volumes was performed and the elution was carried out with 2.5 volumes of elution buffer (50 mM Tris-HCl pH 8.0, 150 mM NaCl, 350 mM Imidazole, 10% glycerol). Finally, imidazole was removed and the protein was concentrated using a Centricon (Merck Millipore) and preserved at −80 °C in small fractions at 2 mM concentration.

### ESI MS for Intact Mass Analysis

The protein samples were analyzed on an LC−ESI-MS platform consisting of a Surveyor MS pump system (C4 column, Higgins Analytical Proto 300 5µm RS-0301-W045, 30 mm × 1.0 mm) coupled to an electrospray mass spectrometer (Thermo Finnigan LCQ duo, equipped with an ion trap mass analyzer). The gradient was 5% solvent B (96% acetonitrile, 2% acetic acid) in solvent A (2% acetonitrile, 2% acetic acid) for 2 min, 5 to 100 % solvent B in solvent A in 4 min, and 100% solvent B for 8 min at a flow rate of 40 μL min-1. The ESI spectra was deconvoluted to the zero-charge domain by the ProMass Deconvolution Software (Novatia) using a standard parameter set: average mass type, mass tolerance 0.02%, minimum tolerance 2 Da and input m/z range 500–2000 units.

### Circular Dichroism Spectroscopy

CD spectra measurements were carried out at 20 °C with a Jasco J-815 spectropolarimeter. Near-UV CD spectra were collected using cells with path lengths 1.0 cm. High tension voltage was registered simultaneously. Data was acquired at a scan speed of 20 nm min^−1^ (five scans were averaged, scans corresponding to buffer solution were averaged and subtracted from the spectra). Values of ellipticity were converted to molar ellipticity.

### Fluorescent Probe

The tetramethylrhodamine (TMR) labeled peptide with sequence Gly-Gly-GLy-Ser-{Lys-(TMR)}was purchased from Genscript. Its mass, as determined by mass spectrometry analysis, and excitation and emission spectra were compatible with the predicted theoretical values. An absorption coefficient of 87500 M^−1^1 cm^−1^ at 560 nm was used to calculate the TMR concentration in solution. To evaluate the transpeptidation efficiency, a spectrum of free TMR was subtracted from the spectrum of protein in order to correct the 280 nm absorbance value of the protein solution.

### Transpeptidation and Multiplicity

The transpeptidation reaction was performed by the Sortase A enzyme. To obtain a product with high multiplicity (6:10, RBD:BLS) 70 and 35 *μ*M of RBD and BLS respectively were used. For lower ratios (approximately 2:10, RBD:BLS), 70 and 350 *μ*M of RBD and BLS respectively were used. The reaction was started by the addition of Sortase A (0.4 *μ*M final concentration from a 2 mM stock solution preserved at −80 °C). The reaction was performed in a 20 mM Tris-HCl, 150 mM NaCl, pH 7.4, 10 mM CaCl_2_ buffer. CaCl_2_ allowed the activation of Sortase A, therefore the reaction was stopped by the addition of 10 mM EDTA. The reaction previously treated or not with 5 mU of endoglycosidase H during 1 h at 37 °C was analyzed by SDS-PAGE and by SEC-HPLC. In order to improve the efficiency and the irreversibility of the transpeptidation reaction, one of the reaction products, the short peptide Gly-His-His-His-His-His-His splitted by Sortase A from the C-terminal of RBD (**Figure 1** for reference), was extracted from the mixture. For this purpose, we performed the reaction in a concentrator of 10 kDa cutoff under centrifugation at 3000 rpm at 4 °C. Control experiments were performed without centrifugation. All samples were kept at 4 °C.

### Hydrodynamic Behavior of the Proteins

Size exclusion chromatography (SEC-HPLC) was carried out using a Superose-6 column (GE Healthcare). The protein concentration was 10-30 μM, a volume of 50 μL was typically injected, and the running buffer was 20 mM Tris-HCl, 100 mM NaCl, 1 mM EDTA, pH 7.0. The experiment was carried out at room temperature (~25 °C) at a 0.5 mL min-1 flow rate. A JASCO HPLC instrument was used. It was equipped with an automatic injector, a quaternary pump and a UV-VIS UV-2075 (the elution was monitored at 280 nm).

### Interaction between Recombinant RBD and ACE2

The interaction between the covalently coupled product RBD-TMR and the ACE2 soluble domain was studied taking advantage of the presence of a (His)_6_ tag in the C-terminus of ACE2, which is not present in RBD-TMR. ACE2 (1.0 μM) and RBD-TMR (1.0 μM) were incubated during 30 min, with gentle agitation in darkness in the presence of NTA-Ni^2+^ in 20 mM sodium phosphate, 150 mM NaCl, pH 7.40. The complex was separated from free RBD-TMR by centrifugation (1 min at maximum speed in a microcentrifuge). The fluorescence after the incubation was measured using an excitation wavelength of 560 nm, the emission monitored between 570 and 670 nm, a 4 nm bandwidth was used for both excitation and emission. Measurements were performed in a JASCO spectrofluorometer equipped with a thermostated cell holder connected to a circulating water bath set at 25 °C. A path length cell of 0.3 cm was used. Additionally, the complex bound to the NTA matrix was eluted by the addition of imidazole and the fluorescence of the supernatant was analyzed in the same fashion.

### Endotoxin Remotion and Detection

Lipopolysaccharide (LPS)^42^ was removed from the protein preparations by an overnight incubation (4 °C) of the proteins with polymyxin B matrix followed by a second incubation (2-3 h) at room temperature. The matrix was previously equilibrated in 20 mM Tris-HCl, 100 mM NaCl, pH 7.0 buffer solution. We measured the LPS content of purified RBD-BLS preparations as previously described^19^. The endotoxin concentration in the protein samples was determined by Cassará Fundation Laboratories. Briefly, all protein samples were brought to a final volume of 200 μL and the LPS content was measured using a commercial kit from Charles River Endosafe (R160), based on the Limulus amebocyte lysate (LAL) assay chromogenic method, with a sensitivity of 0.015 EU/ml. No LPS was detected in the RBD-BLS preparation following endotoxin removal.

### Immunization Protocols

Mice immunization was carried out by experts from the High Level Technological Service from CONICET (STAN No. 4482), under guidelines from the Institutional Committee for the Care and Use of Laboratory Animals (CICUAL). BALB/c mice were obtained from the animal facility of the Faculty of Veterinary Sciences, University of La Plata (Argentina), and housed at the animal facility of the Instituto de Ciencia y Tecnología Dr. César Milstein, Fundación Pablo Cassará. Six groups of five Females mice (6-8 week-old) (n=5) were immunized subcutaneously with the same amount of RBD protein (15 µg) coupled or not to BLS in the presence or absence of 500 µg aluminum hydroxide (Al(OH)_3_). Two different BLS coupling formulations were obtained (**Table 1**): RBD_6.6_/BLS_10_ (high multiplicity, RBD-BLS HM) and RBD_1.5_/BLS_10_ (low multiplicity, RBD-BLS LM). Additionally, uncoupled RBD + BLS formulation (RBD + BLS, **Table 1**) was tested with the same equimolar conditions used for the RBD-BLS HM group. Pre-immune sera also were collected before starting the immunization. Mice received two immunizations with the same dose at days 0 (antigen prime) and 30 (antigen boost). Additional control animals were injected with 500 μg Al(OH)_3_ per mouse with the same immunization schedule.

Blood samples were obtained at 20 days post-second immunization by venipuncture from the facial vein. After coagulation at room temperature for 1-2 h, blood samples were spun in a centrifuge at 3000 rpm/min for 10 min at 4 °C. The upper serum layer was collected and stored at −20°C.

### Identification of Serum Antibody Against Protein RBD in Mice Using an ELISA Assay

The antibody response against RBD and RBD-BLS was measured using a standard ELISA procedure. Briefly, flat-bottom 96-well plates (Thermo Scientific NUNC-MaxiSorp) were coated with recombinant RBD protein produced in *P. pastoris* at a final concentration of 1 μg/ml (100 μl/well) in phosphate-buffered saline (PBS) coating buffer (pH 7.4) at 4 °C overnight. After blocking (8% nonfat dry milk PBS for 2 h at 37 °C) the plates were washed 5 times with PBS containing 0.05% Tween 20 (PBST). Serially diluted mouse sera were incubated at 37 °C for 90 min in PBS containing 1% non-fat dry milk (blocking solution), and plates were subsequently washed with PBST. For total specific IgG determination, IgG horseradish peroxidase (HRP)-conjugated antibody (DAKO P0447) was diluted 1/1000 in blocking solution and added to the wells. After incubation for 1 h at 37 °C, plates were washed fivetimes with PBST and developed with 3,3’,5,5’-tetramethylbiphenyldiamine (TMB) for 15 min. The reaction was stopped with 50 μl/well of 1 M H_2_SO_4_ (stop solution). The absorbance was measured in a microplate reader (Thermo Multiscan FC ELISA) at 450 nm.

The antibody titer was determined as the inverse of the last dilution that was considered positive, with a cut-off value defined as absorbance at 450 nm of 0.20, which was twice as high as that from a pool of normal mouse sera (from 30 unimmunized animals). Statistical significance was evaluated by the Student’s t-test, using a logarithmic transformation of the ELISA titers. Differences were considered significant if p < 0.05.

### Pseudotyped Lentivirus Neutralization Assay

SARS-CoV-2 S-Pseudotyped lentivirus^27,43^ were produced by co-transfection of HEK-293T cells with plasmids bearing a GFP reporter gene (pLB was a gift from Stephan Kissler, Addgene plasmid #11619; http://n2t.net/addgene:11619; RRID:Addgene_11619), a lentivirus backbone (VRC5602, NIH), and the Spike protein (VRC7475_2019-nCoV-S-WT, NIH). Briefly, HEK-293T cells (2×10^7^) were seeded in a 150-mm tissue culture dish in DMEM medium containing 10% FBS. The next day, cells were transfected with 10 μg of VRC5602, 5 μg pLB-GFP, and 3 μg of VRC7475_2019-nCoV-S-WT in OptiMEM medium using PEI in a 1:3 DNA:PEI ratio. Twenty four hours later, transfection efficiency, indicated as GFP fluorescence, was checked under a fluorescent microscope. Pseudotyped lentivirus supernatants were harvested 48 h post-transfection and stored at 4 °C, fresh media (DMEM + 5% FBS) was added to the plates. After 48 h, combined supernatants were clarified by centrifugation for 10 min at 3,000 rpm to pellet the residual cells. The clarified supernatant was centrifuged for 5 h at 10,000 rpm. The pellet was resuspended in a storage medium (OptiMEM + 6% Sucrose) and aliquots were frozen at −80 °C until use.

Pseudotyped lentivirus titers were measured by transducing HEK-293T cells previously seeded in 96-well plates (2×10^4^ cells/well) and transiently transfected with 100 ng of ACE2 (NIH) and 10 ng of TMPRSS2 protease (NIH) per well. The pseudotyped virus stock (concentrated supernatant) was serially diluted in assay medium (DMEM + 2.5% FBS), incubated for 2 h at 37 °C and added to the transfected cells. Virus titers were calculated by counting the GFP positive cells using an automated counting tool in ImageJ (NIH), a titer of 200-225 GFP positive cells/field using 100X magnification was used for the assay.

Neutralization assays were performed on HEK-293T cells transiently transfected 24 h before transduction with ACE2 and TMPRSS2 protease genes. Fifty microliters of serial 5-fold diluted heat-inactivated serum (56 °C for 45 min), starting dilution of 1:100, were prepared in assay medium and incubated for 2 h with an equal volume of titrated pseudotyped lentivirus. Pseudovirus and sera dilution mixtures (100 μl) were then added onto 96-well plates containing 2×10^4^ cells/well. Pseudovirus infectivity was scored 48 h later and images were obtained using an inverted microscope (IX-71 OLYMPUS), and analyzed with the Micro-Manager Open Source Microscopy Software. Sera antibody neutralization titers were calculated by a nonlinear regression curve fit using GraphPad Prism software Inc. (La Jolla, CA). Half maximal inhibitory concentration (IC_50_), corresponding to the serum antibody dilution causing a 50% reduction of GFP positive cells compared to control “virus only” treated cells, was determined using the same software according to Ferrara and Temperton^44^.

## Abbreviations

ACE2: angiotensin-converting enzyme 2
BLS: lumazine synthase from *Brucella abortus*
CD: circular dichroism
CoV: Coronavirus
EDTA: Ethylenediaminetetraacetic acid
ELISA: enzyme-linked immunosorbent assay
ESI-MS: electrospray ionization mass spectrometry
GFP: green fluorescent protein
HM: high multiplicity
HPLC: high-performance liquid chromatography
IPTG: Isopropyl β-D-1-thiogalactopyranoside, Isopropyl β-D-thiogalactoside
LM: low multiplicity
LPS: lipopolysaccharide
LS: lumazine synthase
NTA-Ni2+: nickel-charged nitrilotriacetic acid affinity resin
PBS: phosphate-buffered saline
PDB: Protein Data Bank
PEI: Polyethylenimine
RBD: receptor binding domain
RBD-BLS: the multimeric particle constituted by RBD covalently coupled to the BLS decamer
RBD-TMR: RBD covalently coupled to tetramethylrhodamine
SARS-CoV-1: severe acute respiratory syndrome coronavirus 1
SARS-CoV-2: severe acute respiratory syndrome coronavirus 2
SDS-PAGE: polyacrylamide gel electrophoresis
SEC: size exclusion chromatography
TEV: Tobacco Etch Virus protease
TMR: tetramethylrhodamine
TFA: trifluoroacetic acid
SDS-PAGE: SDS polyacrylamide gel electrophoresis.

## Acknowledgments

We thank LANAIS-PRO-EM for the support with mass spectrometry analysis of proteins and peptides, and Fundación Ciencias Exactas y Naturales from Universidad de Buenos Aires for their help. Also we would like to thank Santigo Sosa for suggestions concerning BLS and Dr. Fernando Goldbaum for providing us a BLS variant without Cys residues. We thank Erik Procko for providing us with pcDNA3-sACE2(WT)-8his (Addgene plasmid #149268), Stephan Kissler for providing us with pLB (Addgene plasmid #11619) and David Liu for providing us with Sortase A pentamutant (Addgene plasmid #75144).

## Author Contributions

All authors from the Argentinian AntiCovid Consortium (*) designed experiments, performed research, analyzed data and drafted the manuscript. All authors (listed in alphabetical order) contributed equally to this work. P. C. F. determined endotoxin contents in the protein samples prior to the formulation for immunizations.

## UniProt Accession IDs

UniProtKB - P0DTC2 (SPIKE_SARS2); Spike glycoprotein from SARS-CoV-2.

UniProtKB - Q2YKV1 (RISB2_BRUA2); 6,7-dimethyl-8-ribityllumazine synthase 2.

UniProtKB - Q9BYF1 (ACE2_HUMAN); human angiotensin-converting enzyme 2.

