## Supplementary Material for "Production of a Highly Immunogenic Antigen from SARS-CoV-2 by Covalent Coupling of the Receptor Binding Domain of Spike Protein to a Multimeric Carrier"

***Sequences of proteins used in this work.***

Sortase A pentamutant (eSrtA):

MQAKPQIPKDKSKVAGYIEIPDADIKEPVYPGPATREQLNRGVSFAEENESLDDQNISIAG
HTFIDRPNYQFTNLKAAKKGSMVYFKVGNETRKYKMTSIRNVKPTAVEVLDEQKGKDK
QLTLITCDDYNEETGVWETRKIFVATEVKLEHHHHHH

(yellow, C-terminal Histag)

BLS sequence (modified for coupling by Sortase A enzyme):

MHMENLYFQGGGGSGSGLKTSFKIAFIQARWHADIVDEARKSFVAELAAKTGGSVEVEI
FDVPGAYEIPLHAKTLARTGRYAAIVGAAFVIDGGIYRHDFVATAVINGMMQVQLETEVP
VLSVVLTPHHFHESKEHHDFFHAHFKVKGVEAAHAALQIVSERSRIAALV

(in green, the TEV site and in orange a Sortase A site (N-terminal Gly-Gly-Gly) and a short linker Ser-Gly-Ser-Gly)

ACE2 Soluble receptor domain sequence:

MSSSSWLLLSLVAVTAAQSTIEEQAKTFLDKFNHEAEDLFYQSSLASWNYNTNITEENVQ
NMNNAGDKWSAFLKEQSTLAQMYPLQEIQNLTVKLQLQALQQNGSSVLSEDKSKRLN
TILNTMSTIYSTGKVCNPDNPQECLLLEPGLNEIMANSLDYNERLWAWESWRSEVGKQL
RPLYEEYVVLKNEMARANHYEDYGDYWRGDYEVNGVDGYDYSRGQLIEDVEHTFEEI
KPLYEHLHAYVRAKLMNAYPSYISPIGCLPAHLLGDMWGRFWTNLYSLTVPFGQKPNID
VTDAMVDQAWDAQRIFKEAEKFFVSVGLPNMTQGFWENSMLTDPGNVQKAVCHPTAW
DLGKGDFRILMCTKVTMDDFLTAHHEMGHIQYDMAYAAQPFLLRNGANEGFHEAVGEI
MSLSAATPKHLKSIGLLSPDFQEDNETEINFLLKQALTIVGTLPFTYMLEKWRWMVFKGE
IPKDQWMKKWWEMKREIVGVVEPVPHDETYCDPASLFHVSNDYSFIRYYTRTLYQFQF
QEALCQAAKHEGPLHKCDISNSTEAGQKLFNMLRLGKSEPWTLALENVVGAKNMNVR
PLLNYFEPLFTWLKDQNKNSFVGWSTDWSPYADGSGHHHHHHHH

(green, the natural signal peptide of ACE2 and in yellow, a short linker Gly-Ser-Gly and the C-terminal His tag)

**RBD sequence for *Pichia pastoris* expression:**

MRFPSIFTAVLFAASSALAAPVNTTTEDETAQIPAEAVIGYSDLEGDFDVAVLPFSNSTNNG
LLFINTTIASIAAKEEGVSLEKREAEAEFRVQPTESIVRFPNITNLCPFGEVFNATRFASVYA
WNRKRISNCVADYSVLYNSASFSTFKCYGVSPTKLNDLCFTNVYADSFVIRGDEVRQIAP
GQTGKIADYNYKLPDDFTGCVIAWNSNNLDSKVGGNYNYLYRLFRKSNLKPFERDISTE
IYQAGSTPCNGVEGFNCYFPLQSYGFQPTNGVGYQPYRVVVLSFELLHAPATVCGPKKS
TNLVKNKLPETGHHHHHH

(green alpha factor, yellow Sortase A and C-terminal Histag sequences)

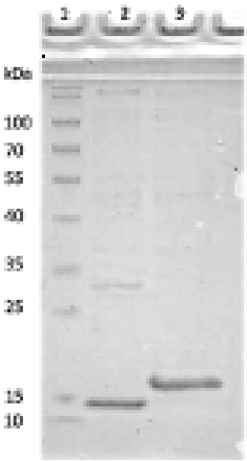


**Figure S1. SDS-PAGE Analysis of BLS Digestion by TEV Protease.** Lane 1: molecular weight markers, lane 2: BLS treated with TEV protease and lane 3: intact BLS. BLS was purified by ion exchange chromatography followed by SEC-HPLC**.**

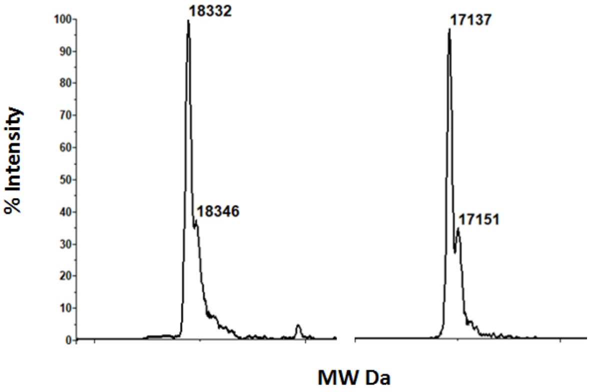


**Figure S2. ESI-MS Spectra of BLS.** Spectra corresponding to *Met-Glu-Asn-Leu-Tyr-Phe-Gln*-Gly-Gly-Gly-**BLS** (left) and Gly-Gly-GLy-**BLS**, after digestion with TEV protease (right) are shown.

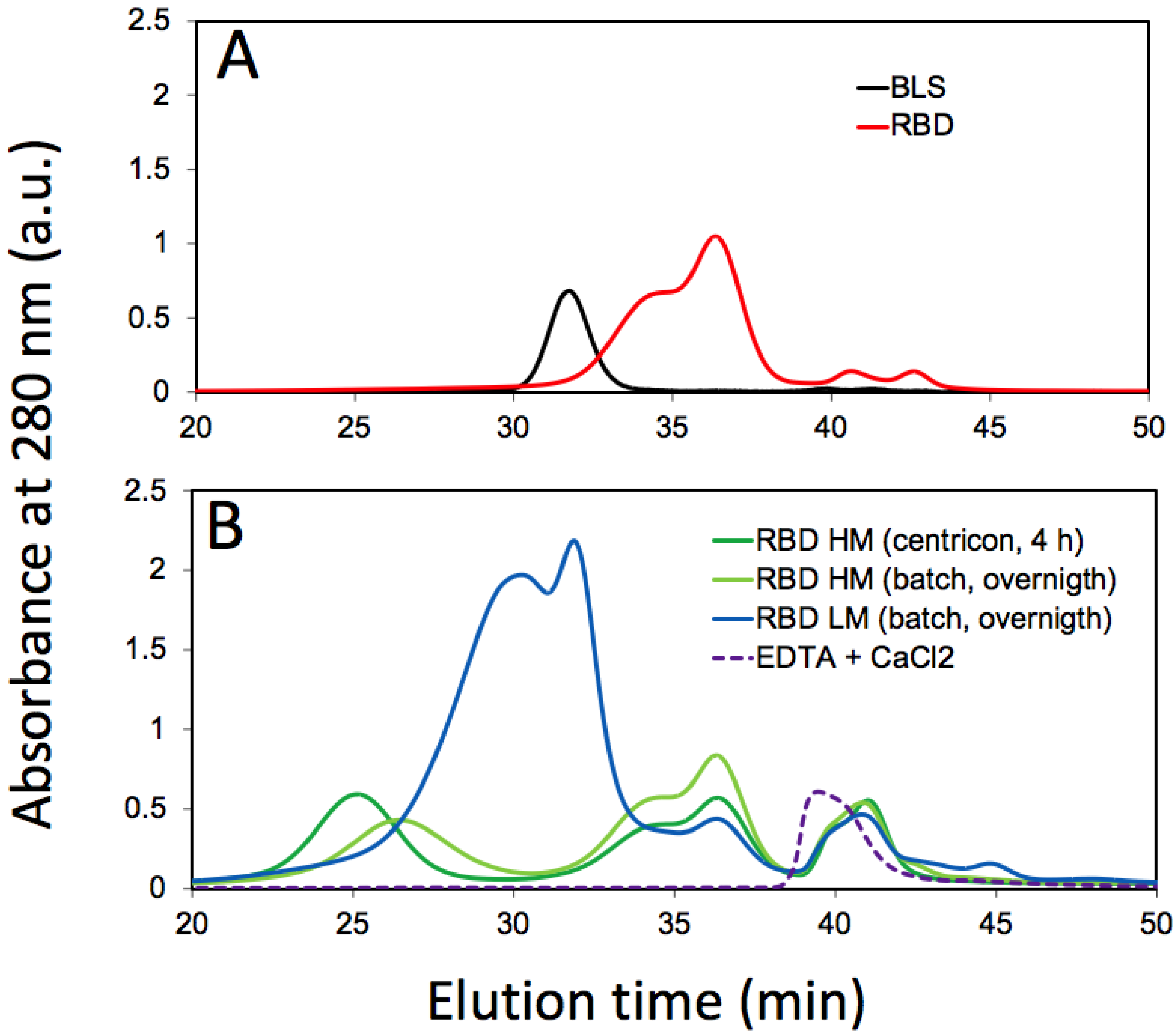


**Figure. S3. Sortase A-mediated covalent coupling of RBD and BLS at low and high multiplicities.** (A) SEC-HPLC profiles corresponding to Gly-Gly-Gly-BLS (black), RBD (red). (B) The product of the reaction catalyzed by Sortase A prepared under two different conditions: high multiplicity and Centricon 4h (dark green) and batch overnight (light green) or prepared at low multiplicity (blue) . An equivalent volume of a 10 mM EDTA, 10 mM $CaCl_2$ solution was loaded as a control (magenta, lower pane). Decameric BLS has a molecular weight of approximately 170 kDa and each RBD subunit adds approximately 26 kDa or 40 kDa (excluding or including glycosylation, respectively). The peak. observed between 38-43 min corresponds to EDTA-$Ca^{2+}$.

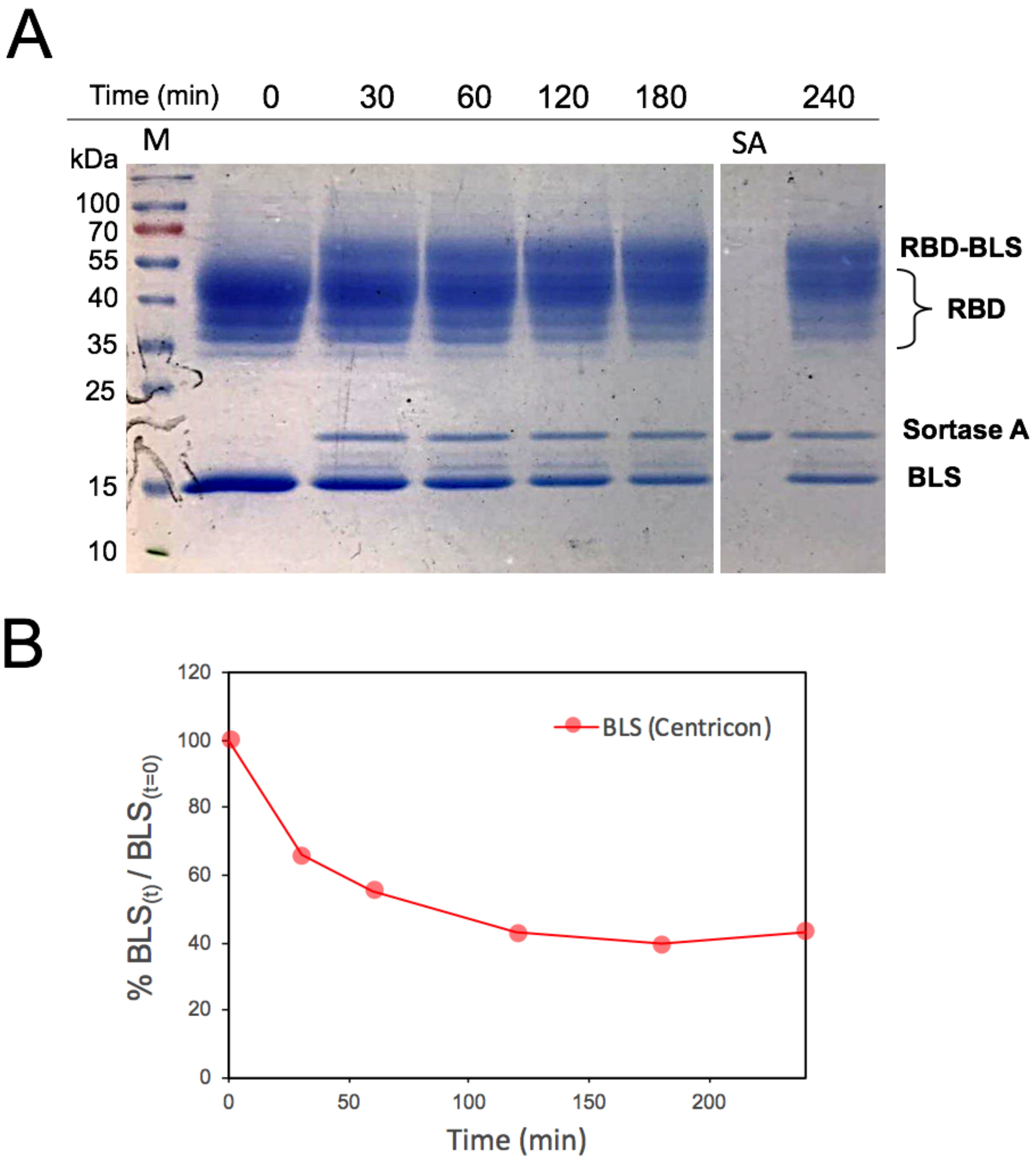


**Figure S4. Monitoring RBD-BLS Covalent Coupling.** (A) Sortase A reaction was monitored by an SDS-PAGE analysis. The reaction was carried out at 4 ºC in a centrifugal filter unit (Centricon), as described in Materials and Methods. Small samples were separated at reaction times of 0, 30, 60, 120, 180 and 240 min. Sortase A (SA) was included as a control, and molecular weight markers (M) were included as a reference. (B) The quantification of the loss of BLS band in SDS-PAGE. The ratio BLS$_{(t)}$ /BLS$_{(t=0)}$ was calculated for each reaction time and was plotted as percentage. BLS$_{(t)}$ and BLS$_{(t=0)}$ are the initial and the remaining BLS masses estimated from the analysis of the band density corresponding to the BLS band.
